# Chemoenzymatic fluorescence labeling of intercellularly contacting cells using lipidated sortase A

**DOI:** 10.1101/2022.05.09.491123

**Authors:** Satoshi Yamaguchi, Ryosuke Ikeda, Yuki Umeda, Takahiro Kosaka, Shinya Yamahira, Akimitsu Okamoto

**Author notes:** Corresponding author: Satoshi Yamaguchi, Department of Chemistry & Biotechnology, School of Engineering, The University of Tokyo, 7-3-1 Hongo, Bunkyo-ku, Tokyo 113-8656, Japan, /Fax, Akimitsu Okamoto, Department of Chemistry & Biotechnology, School of Engineering, The University of Tokyo, 7-3-1 Hongo, Bunkyo-ku, Tokyo 113-8656, Japan; RCAST, The University of Tokyo, 4-6-1 Komaba, Meguro–ku, Tokyo 153-8904, Japan, /Fax.

## Abstract

Methods to label intercellular contact attract particular attention due to their potential in cell biological and medical applications through analysis of intercellular communications. In this study, a simple and versatile method for chemoenzymatically labeling the intercellularly contacting cell was developed by using a cell-surface anchoring reagent of poly(ethylene glycol)(PEG)-lipid conjugate. The surfaces of each cell in cell pairs of interest were efficiently decorated with sortase A (SrtA) and triglycine peptide that were lipidated with PEG-lipid, respectively. In the mixture of the two cell populations, the triglycine-modified cells were enzymatically labeled with a fluorescent labeling reagent by contacting with the SrtA-modified cells both on the substrate and in cell suspensions. Such selective labeling of the contacting cells was confirmed by confocal microscopy and flow cytometry. The results show a proof of principle that the present method is a promising tool for selective visualization and quantification of the intercellularly contacting cells among cell mixtures in cell-cell communication analysis.

## Introduction

Cell-cell communications via the intercellular membrane contact are critical for regulating cellular functions in a variety of biological phenomena [1,2]. Therefore, detection and quantification of cell-cell interactions are important for giving new insights into fundamental cell biology [3,4], diagnosing the activity of disease-related cells [5] and improving quality control of therapeutic cells [6]. Until now, cell-cell interactions also have been examined simply by mixing multiple cell populations, in which heterogeneous cell pairs happened to interact with each other. Recently, techniques have been actively developed for quantitatively observing the interactions of single-cell pairs to comprehensively investigate the heterogeneity of cell-cell communications [7-10]. However, these techniques only measured the cellular responses presumably induced by cell-cell interactions without directly examining the intercellular contacts. Accordingly, it is not strictly known whether such observed responses were caused by cell-cell interactions or not.

In such context, fluorescence labeling techniques to detect intercellular contacts have been reported extensively in recent years. A proximity-dependent enzymatic labeling method using the *Staphylococcus aureus* transpeptidase sortase A (SrtA) has been developed for detection of intercellular molecular interactions [11,12]. Here, SrtA has been extensively employed for site-specific linkage of proteins and peptides *in vitro* or on living cells in various applications [13,14], particularly for protein modification with synthetic functional molecules such as lipids [15,16] and fluorescent dyes [17,18]. In such SrtA-mediated transpeptidation, SrtA recognizes an LPXTG sequence, where X is a variable amino acid and cleaves the amide bond between T and G, resulting in the formation of the thioester intermediate with the carboxyl group of T at the active-site cysteine. This thioester intermediate undergoes nucleophilic attack by an *N*-terminal amine group of the oligoglycine sequence, thereby ligating two peptides including LPXTG and oligoglycine sequences. In the pioneering method for labeling intercellular contact, two populations of cells were transfected separately with genes for expressing membrane proteins fused with SrtA and oligoglycine, respectively. The two cell populations were mixed in the presence of a fluorescence-labeled LPETGG peptide as the labeling agent. At sites in close enough proximity to allow intermolecular contact between these heterogeneous cells, SrtA on the cell membrane can link the labeling agent to the oligoglycine on the neighboring partner cell. Using this technique, fluorescence labeling of the contact sites between immune cells has been reported [11]. Alternatively, the *N*- and *C*-terminal domains of a split green fluorescent protein (split GFP) were reported to be expressed on the cell membranes of heterogenous cells using glycosylphosphatidylinositol (GPI) anchor, and the intercellular contacting cells were fluorescently labeled through proximity-based reconstruction of split GFP [19]. Thus, techniques to visualize cell-cell contacts by fluorescence microscopy have been actively reported in recent years. However, these techniques require the introduction of each protein gene into the heterogenous cell pair of interest. Therefore, they are time-consuming and difficult to be applied to cells with low gene transfer efficiency.

In this study, we develop a simple and versatile method for chemoenzymatic fluorescence labeling of intercellularly contacting cells without using gene transfer. SrtA and oligoglycine peptides were introduced onto the cell membrane using a cell-surface anchoring reagent consisting of poly(ethylene glycol)(PEG) and lipid. Here, PEG-lipids are spontaneously incorporated into cell membranes from extracellular regions via the interaction of their lipid moiety with lipid bilayer membranes. Proteins and peptides have been reported to be modified onto live-cell surfaces through their lipidation by PEG-lipid [20-22]. In particular, a PEG conjugate with dioleoyl phosphatidylethanolamine (PEG-DOPE) achieved stable protein modification of cell surfaces for a relatively long period in culture medium [20]. Therefore, SrtA and a triglycine-including peptide were chemically lipidated with PEG-DOPE (Fig. 1A), and the lipidated SrtA (SrtA-lipid) and the lipidated peptide (G3-lipid) were introduced onto each cell surface of heterogenous cell pairs. SrtA-lipid modified and G3-lipid modified cells were mixed under the existence of fluorescently-labeled LPETGG-including peptide, and the fluorescence labeling through proximity-dependent ligation was detected from the cells with the intercellular contact (Fig. 1B).

**Fig. 1.**
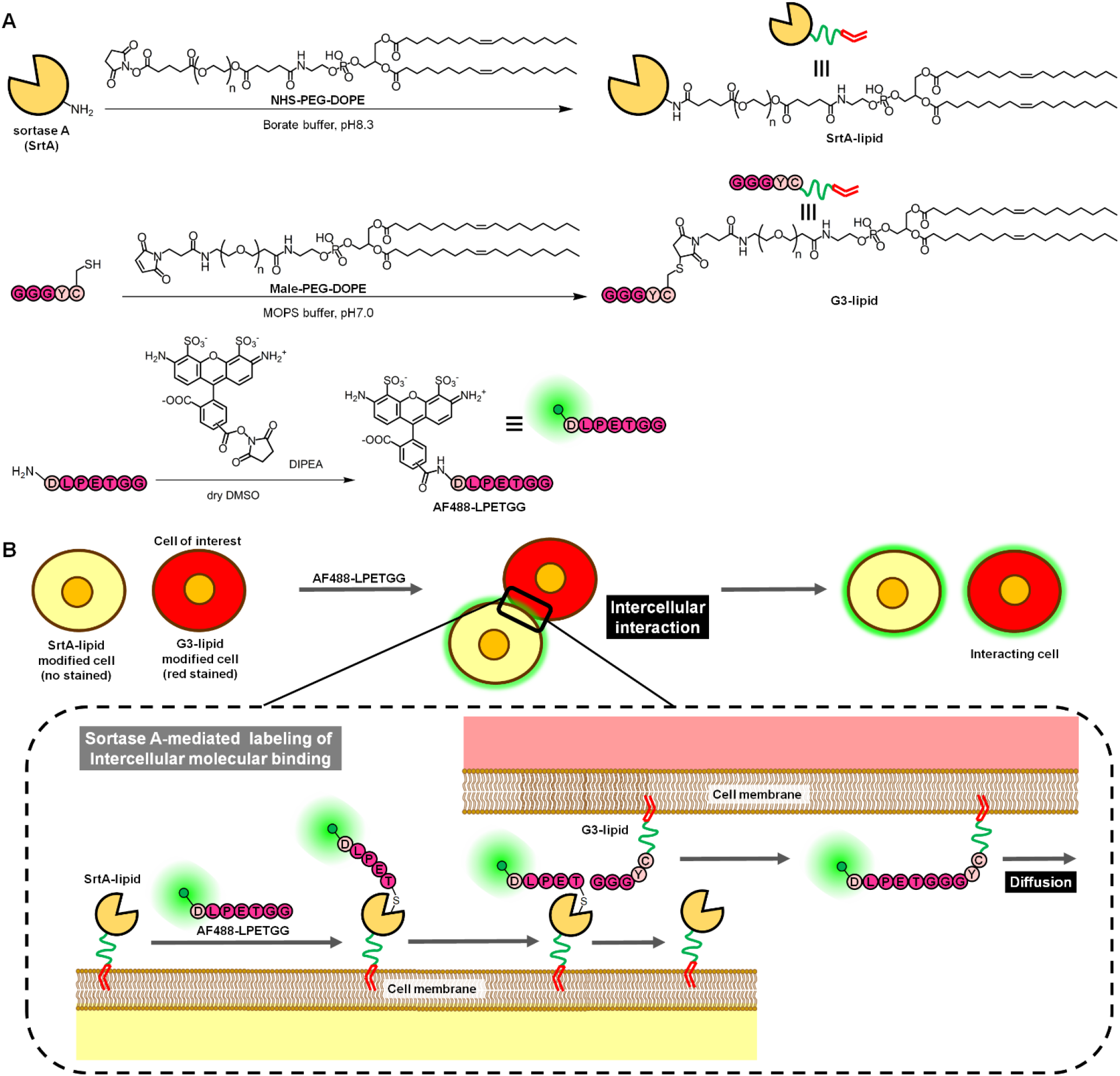
(A) Chemical structures of lipidated enzyme and peptide and a labeling regent. Lipidated Sortase A (SrtA-lipid) and lipidated triglycine-including peptide (G3-lipid) were prepared by conjugation with NHS-PEG-DOPE and Male-PEG-DOPE, respectively. AF488-labeled LPETGG-including peptide (AF488-LPETGG) was synthesized as a green fluorescence labeling reagent. (B) Schematic of fluorescence-labeling of the interacting cell of interest. In the sortase A-mediated labeling, fluorescently labeled cells of interest and their interacting partner cells were modified with G3-lipid and SrtA-lipid, respectively, and then, mixed in AF488-LPETGG-including media, resulting in labeling of the cells of interest with green fluorescence.

## Materials and methods

### Materials

PEG (Mn: 5000 (g/mol)) disuccinimidyl ester (NHS-PEG-NHS) (Sunbright DE-050GS), DOPE (Coatsome ME-8181) and oleoyl-PEG-disuccinimidyl ester (NHS-PEG-lipid) (Sunbright OE-040CS) were purchased from NOF Co. (Tokyo, Japan). Alexa Fluor™ 488 succinimidyl ester (AF488-NHS) and Alexa Fluor™ 647 succinimidyl ester (AF647-NHS) were from Thermo Fisher Scientific Inc. (Waltham, MA, USA). CytoRed was from Dojindo Laboratories (Kumamoto, Japan). RPMI1640 medium was from FUJIFILM Wako Pure Chemical Corporation. Penicillin-Streptomycin Mixed Solution and accutase were from Nacalai Tesque, Inc. Fetal bovine serum (FBS) was from Biowest (Nuaillé, France). Type I collagen solution (3.0 mg/mL, Cell Matrix Type I-A) was from Nitta Gelatin (Osaka, Japan). PEG-DOPE succinimidyl ester (NHS-PEG-DOPE) was synthesized by mixing NHS-PEG-NHS (185 mg, 37 μmol) and DOPE (24.7 mg, 33 μmol) with existence of triethylamine (50 μl, 360 μmol) in 10 ml of anhydrous CH_2_Cl_2_. NHS-PEG-DOPE was purified by precipitation in diethyl ether and dried to be obtained as a white solid. G3-lipid was prepared by conjugation of GGGYC peptide with maleimidyl PEG (Mn: 4000 (g/mol))-DOPE by following our previous report [16]. AF488-labeled DLEPTGG peptide (AF488-pep) was prepared by mixing DLEPTGG peptide with AF488-NHS as described in our previous report [23].

### Lipidation and fluorescence labeling of SrtA

SrtA was expressed and purified by following our previous report [16]. SrtA, NHS-PEG-DOPE and AF647-NHS (with the final concentrations of 100, 500 and 100 μM) were mixed in sodium borate buffer (100 mM, pH 8.3). After incubation at room temperature for 1 hour, the conjugation reaction was stopped by adding a small portion of Tris-HCl buffer (1 M, pH8.0). The AF647 labeled SrtA-lipid (AF647-SrtA-L) was purified by dialysis into phosphate-buffered saline (PBS), and then, characterized by SDS-PAGE analysis with CBB staining. The mean AF647-labeling ratio to one SrtA molecule was determined to be 0.6 by the UV/vis absorbance measurement with NanoDrop1000 spectrometer (Thermo Scientific, Willington, USA).

### SrtA modification of cells

Human T cell leukemia (Jurkat) was Riken Bioresource research center (Ibaraki, Japan) and cultured in RPMI1640 medium supplemented with 10 v/v% FBS, 100 u/mL penicillin, and 100 μg/mL streptomycin. Cells were washed with PBS twice and then, suspended in a solution of AF647-SrtA-L (50 μM in PBS) for 15 min at room temperature. After centrifugation and replacement of the solution with PBS twice, the cells suspension was put on the glass-bottom dish and observed with a confocal laser scanning microscope (TCS SP8, Leica Microsystems GmbH, Germany).

### Confocal microscopic observation of intercellular contact

SrtA-modified cell (SrtA-Cell) and triglycine-modified cell (G3-Cell) were prepared by treating Jurkat cells with AF647-SrtA-L and G3-lipid, respectively. A suspension of Jurkat cell (4 × 10^5^ cells/ml) in PBS was divided to two portions, and one portion was stained in red by incubation in a CytoRed solution (2 nM in PBS) for 15 min. The non-stained cells were modified with AF647-SrtA-L as described above. Similarly, the red-stained cells were modified with G3-lipid (50 μM in PBS). Here, the G3 peptide was reported to modify the cell membrane with the same G3-lipid in our previous report [16]. These SrtA-Cell and G3-Cell suspension (each 4 × 10^3^ or 1.2 × 10^6^ cells/ml in PBS) were mixed in a microtube. The glass-bottom dish was coated with collagen by immersing overnight in a solution of collagen (0.3 mg/mL in deionized water at pH3). The collagen surface was immersed in an NHS-PEG-lipid solution (100 μM in PBS) for 1 hour. The cell mixture suspensions were put onto the PEG-lipid-modified surface for cell anchoring on the dish. After incubation for 30 min, the dish surface was rinsed with PBS, and then, the immobilized cell mixture was incubated in a SrtA-mediated labeling solution (40 μM AF488-pep, 50 mM Tris, 150 mM NaCl, 5 mM CaCl2, pH 8.0) for 80 min. After rinsing the dish surface with PBS twice, the cell mixture was observed by confocal microscopy as described above. As negative control, the similar experiments were performed without using AF647-SrtA-L, G3-lipid and/or AF488-pep, respectively.

### Flow cytometry analysis of intercellular contact

The suspensions of SrtA-Cell and G3-Cell (each 6 × 10^7^ cells/ml in PBS) were prepared as described above. In this experiment, G3-Cell was not stained in red because the fluorescence of AF647 and CytoRed could not be distinguished under the experimental setting of flow cytometry. These modified cells were mixed in a microtube. After centrifugation, the supernatant was replaced with the SrtA-mediated labeling solution, and the cell mixtures were resuspended in the solution, followed by incubation for 1 hour at room temperature. After centrifugation and replacement of the supernatant with PBS twice, the cell mixtures were analyzed with a flow cytometer (Guava easyCyte 8, from Luminex Japan Co. Ltd., Tokyo, Japan). As negative control, the similar experiments were performed without using AF647-SrtA-L, G3-lipid and/or AF488-pep, respectively. For quantification of the contacting cells, the AF488-fluorescence positive cell was counted among the AF647-fluorescence negative cell fraction by using FlowJo software (Treestar Inc, San Carlos, CA). The threshold for AF488-fluorescence positivity was determined to set the positive rate of the negative control one without using G3-lipid to 5%.

## Results and discussion

### Cell surface modification with SrtA

SrtA was first modified onto the cell membrane without gene transfer. As mentioned above, SrtA was lipidated via conjugation with PEG-DOPE and simultaneously, labeled with a fluorescent dye, AF647 for visualization on the cell (Fig. 1A). In SDS-PAGE analysis, the ladder bands of the lipidated and labeled SrtA (AF647-SrtA-L) were observed at the upstream region compared with that of intact SrtA (Fig. 1B, closed triangle). These three bands were suggested to be derived from SrtA with various modification numbers of PEG-DOPE from one to three. By quantification with image analysis of the bands, the modification number was mainly one or two, and total 73 % of SrtA was lipidated. In confocal microscopic observation, the AF647 fluorescence was observed from the surface of the Jurkat cells treated with AF647-SrtA-L (Fig. 1C), whereas such fluorescence was not observed from non-treated cells (Fig. 1D). This result clearly shows that SrtA were modified onto cell surfaces by treatment of cells with AF647-SrtA-L. Furthermore, as shown in Fig. 1C, almost all Jurkat cells were modified with SrtA. It is well-know that the efficacy of gene transfer into most floating cells are low (generally up to almost 30%) in conventional methods such as lipofection [24] and virus infection [25]. Jurkat cell is one of the floating cells. In the present method, almost all Jurkat cell was uniformly modified by simple and rapid treatment with lipidated SrtA.

### Microscopic observation of intercellular contact

We first microscopically observed the mixture of SrtA-modified and G3-modified Jurkat cells (SrtA-Cell and G3-Cell) at a low cell density (Fig. 3). To distinguish the contacting cell pairs among the cell mixture, the floating cells were anchored onto the substrate surface by using a cell-anchoring reagent (Fig. 3A). In these confocal microscopic images, SrtA-Cell was identified from the AF647-fluorescence (*blue*) of AF647-SrtA-L at the cell surfaces (Fig 3E). G3-Cell was identified from the CytoRed fluorescence (*red*) at the cytosol (Fig. 3D). After incubation with the green fluorescence-labeled LPETGG peptide (AF488-pep), the AF488-fluorescence (*green*) was observed at the cell surface of all SrtA-Cell (Fig. 3B). It is reasonably explained that the thioester intermediate with the cysteine of the active site of SrtA was formed on the cells. More importantly, the AF488-fluorescence was also observed at the surface of G3-Cell that seems to contact with SrtA-Cell (Fig. 3B and 3F, white arrow). On the non-contacting other G3-Cell, the AF488-fluorescence was not observed. This result indicates that the AF488-fluorescence on G3-Cell depends on the intercellular contact with SrtA-Cell. In the control experiment without using G3-lipid, the red-fluorescent cells have no G3 peptide at their surface (Fig. 3G-K). In this case, the AF488-fluorescence was not observed at the surface of the red fluorescent cells contacting with SrtA-Cell (Fig. 3G and 3K, white arrow). From these results, the present AF488-fluorescence on G3-Cell contacting SrtA-Cell was strongly suggested to be derived from the ligation of AF488-pep with the G3 peptide at the intercellular contacting site. Here, the AF488-fluorescence was observed at the whole surface of G3-Cell. This result suggests that the AF488-pep-ligated G3-lipids diffused on the cell membrane from the intercellularly contacting site to the whole surface. From the properties of lipidated molecules, it is reasonable, and the similar diffusion to the whole cell membrane was reported in the reported method using GPI-anchored split GFP [19]. Thus, the intercellularly contacting cells were confirmed to be labeled with the fluorescent dye simply by treatment with the PEG-lipid-conjugated SrtA and G3 peptide.

**Fig. 2.**
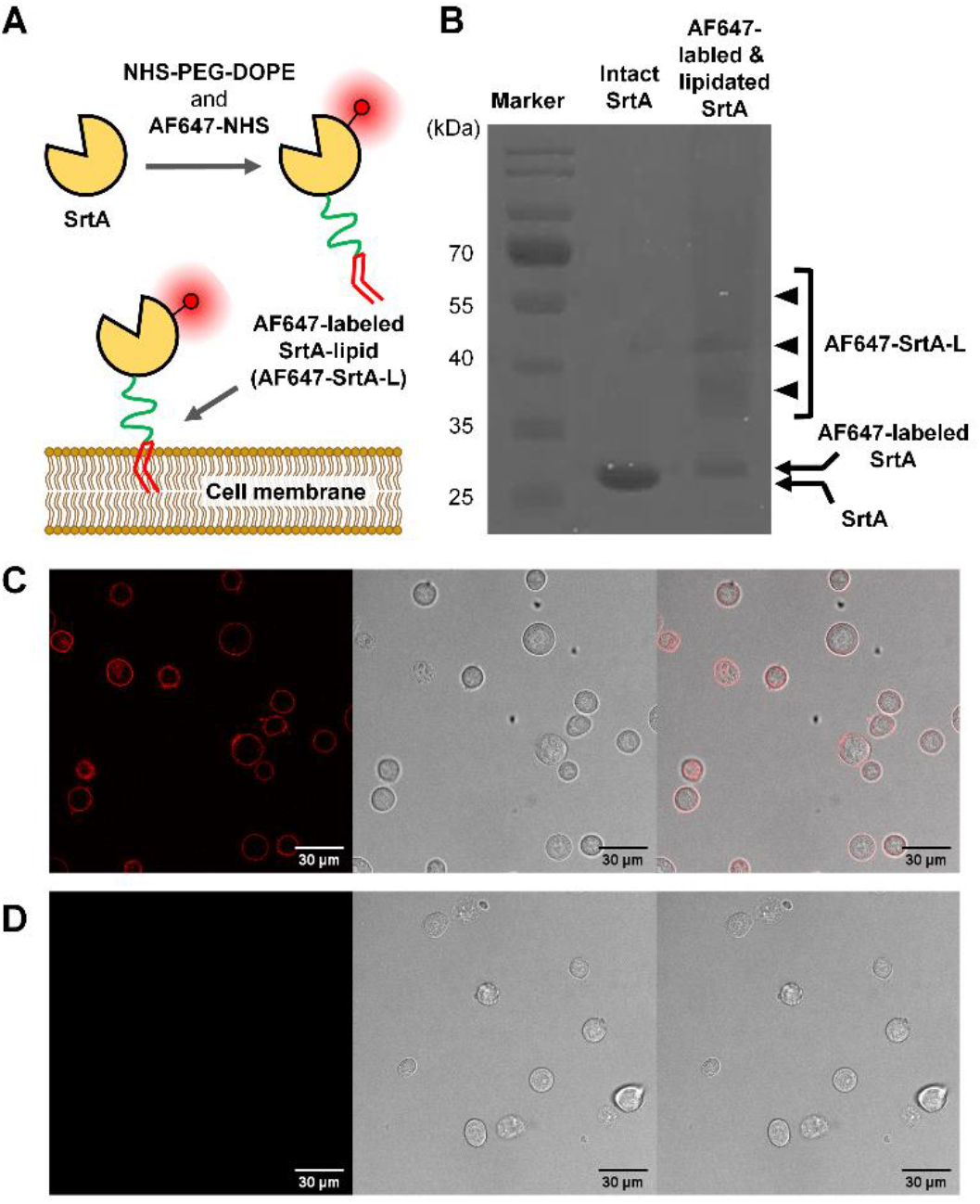
Anchoring of sortase A onto cell membranes through lipidation. (A) Schematic of lipidation and fluorescence (AF647)-labeling of sortase A (SrtA) and its anchoring onto cell membranes. (B) Photograph of SDS-PAGE gel of SrtA treated with and without a fluorescence labeling reagent (AF647-NHS) and a lipidation reagent (NHS-PEG-DOPE). (C, D) Confocal fluorescence microscopic image of Jurkat cells treated (C) with and (D) without AF647-labeled SrtA-lipid (AF647-SrtA-L). (*left*)The AF647 fluorescence image in red, (*center*) the differential interference contrast one and (*right*) the merged one were represented. Scale bar: 30 μm.

**Fig. 3.**
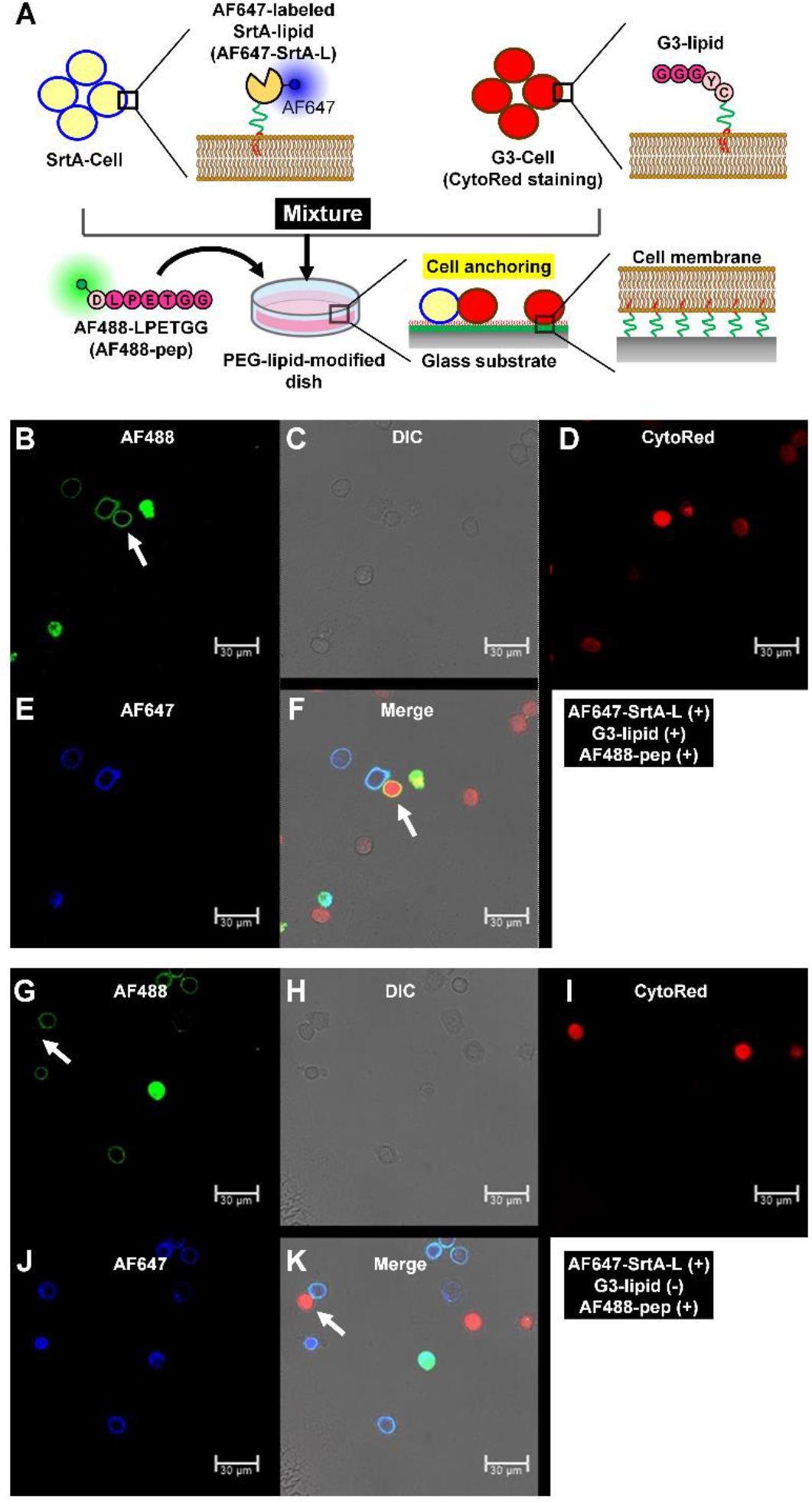
Microscopic observation of fluorescence labeling of interacting cells. (A) Schematic of sortase A (SrtA)-mediated labeling of interacting cells on the substrate. SrtA-Cell and G3-Cell were prepared by treatment of Jurkat cells with AF647-labeled SrtA-lipid (AF647-SrtA-L) and G3-lipid, respectively. G3-Cell was stained in red with CytoRed. These cells were mixed and anchored on the substrate, and then, the interacting ones of G3-Cell were green fluorescently labeled through ligation with AF488-LPETGG peptide (AF488-pep). (B-F) Confocal microscopic images of the mixture of SrtA-Cell and G3-Cell after incubation with AF488-pep. (B) The AF488-fluorescence image in green, (C) the differential interference contrast (DIC) one, (D) the CytoRed-fluorescence one in red, (E) the AF647-fluorescence one in blue and (F) the merged one were observed. The white arrow indicates the interacting G3-Cell. (G-K) The control images were observed without G3-lipid. The images of (G), (H), (I), (J) and (K) were observed in the same way as (B), (C), (D), (E) and (F), respectively. The white arrow indicates the interacting non-treated cell.

Next, the intercellular contact of SrtA-Cell and G3-Cell was observed at a high cell density (Fig. 4). In the high-density cell mixture, the AF488-fluorescence was observed from the G3-Cell contacting with SrtA-Cell, similar to the low-density one (Fig. 4A and 4E). With more careful observation, the AF488-fluorescence was also observed from the G3-Cell that seemed not to contact with SrtA-Cell in the confocal image (typically, the cell indicated with the white ‘X’ in Fig. 4E). But, when the focal point (the z position) of the confocal image was set near the bottom of the dish, the indicated G3-Cell of interest was clearly confirmed to contact with a few SrtA-Cell at the sites near the bottom (Fig. 4F). Different from the present images, in the control experiments without AF647-SrtA-L, G3-lipid and/or AF488-pep, no AF488-fluorescence of the ring-shape was observed from the surfaces of G3-Cell regardless of the intercellular contact (Fig. 4G-4J). From these results, the intercellularly contacting cells were confirmed to be selectively labeled even at a high cell density.

**Fig. 4.**
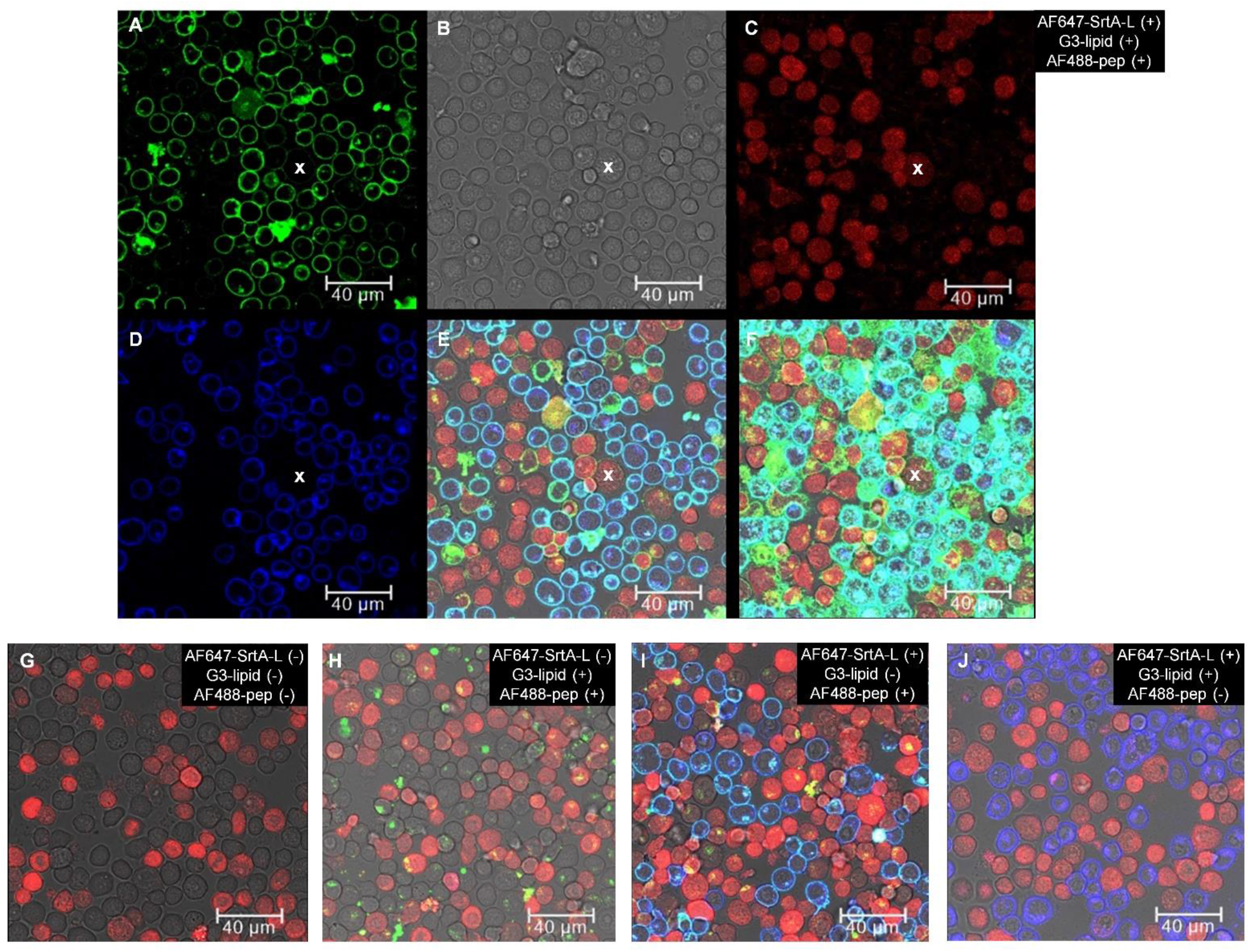
Microscopic observation of fluorescence labeling of interacting cells at high cell density. (A-F) Confocal microscopic images of the mixture of SrtA-Cell and G3-Cell after incubation with AF488-LPETGG peptide (AF488-pep). The images of (A), (B), (C), (D) and (E) were observed in the same way as (B), (C), (D), (E) and (F) in Fig.3, respectively. The image of (F) was observed at the same x-y position as (E) and at the lower z position near the bottom. The white ‘X’ indicates a typical G3-Cell that interacted with SrtA-Cell only near the bottom. (G-J) The control images were observed (G) without AF647-SrtA-L, G3-lipid and AF488-pep, (H) without AF647-SrtA-L, (I) without G3-lipid and (J) without AF488-pep.

### Flow cytometry analysis of intercellular contact

Finally, the intercellular contact of G3-Cell with SrtA-Cell was quantitatively analyzed by flow cytometry. Before flow cytometry analysis, the mixture of SrtA-Cell and G3-Cell was incubated at the bottom of the microtube for 1 hour under the existence of AF488-pep for labeling the contacting G3-Cell (Fig. 5A). The AF488-positive ratio of G3-Cell was 19%, whereas that in the control experiment without G3-lipid was 5.0% (Fig. 5B). This result indicates that the fluorescence increase derived from SrtA-mediated intercellular labeling could be measured by flow cytometry. In another control experiment without AF-pep, the positive ratio was 0.22% (Fig. 5B, light blue). From these results, the fluorescence of non-specifically adsorbed AF-pep onto the cells is estimated to be almost 5.0% and is included in the positive ratio in the experiments using AF-pep. Here, in the control experiment using free SrtA in the mixture of non-treated cells and G3-Cell, the positive ratio was 56% (Fig. 5C). In this positive control, all G3-Cell is set to be fluorescently labeled by free SrtA. Considering the non-specific fluorescence of almost 5%, this positive ratio (56%) indicates that all G3-Cell was reasonably detected as the positive cell. Similarly, by subtracting the non-specific fluorescence, the intercellularly contacting G3-Cell with SrtA-Cell in the microtube was assumed to be approximately one-fifth of G3-Cell. Thus, the present labeling method is also potentially available for quantifying the intercellularly contacting cells of interest in cell suspensions.

**Fig. 5.**
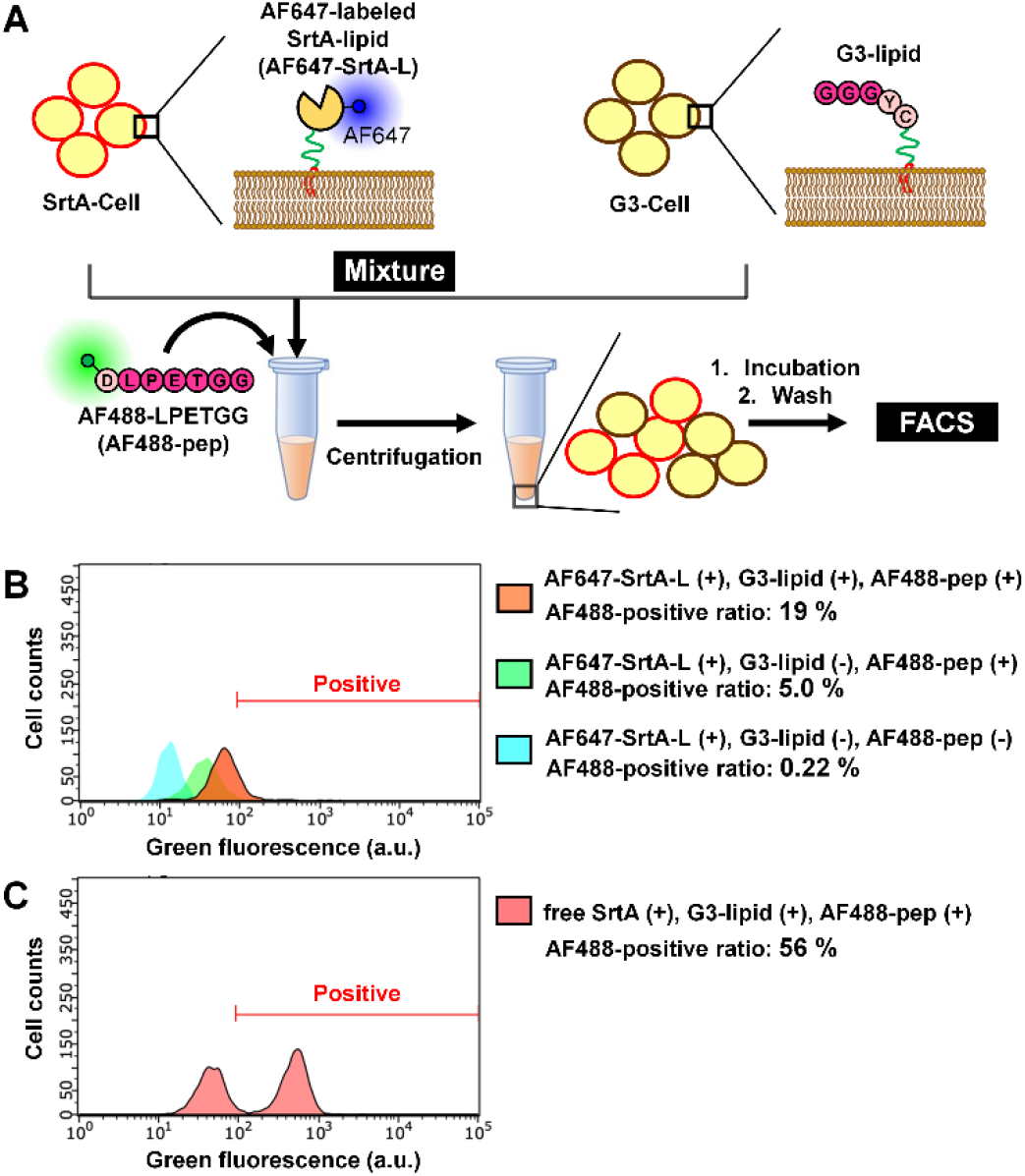
Flow cytometry analysis of fluorescently labeled interacting cells. (A) Schematic of sortase A (SrtA)-mediated labeling of interacting cells in the cell mixture pellet. SrtA-Cell and G3-Cell were prepared by treatment of Jurkat cells with AF647-labeled SrtA-lipid (SrtA-L) and G3-lipid, respectively, and then, mixed and centrifugated. In the pellet at the bottom of microtubes, the interacting ones of G3-Cell were green fluorescently labeled through ligation with AF488-LPETGG peptide (AF488-pep). (B,C) The histograms of green fluorescence intensity of the cells. (B) G3-Cell in the mixture with SrtA-Cell was analyzed with or without G3-lipid and AF488-pep. (C) G3-Cell in the mixture with non-treated Jurkat cells in the existence of free SrtA and AF488-pep was analyzed as a control.

## Conclusions

Simple modification of cells with chemically lipidated SrtA and G3 peptide achieved proximity-based fluorescence labeling of intercellularly contacting cells among their cell mixtures. By using the lipidated SrtA and G3 peptide with PEG-lipid, almost all cells in each cell population could be modified with SrtA and G3, respectively. Since the modification was achieved to be highly efficient even on floating cells, the present method is considered to be more versatile compared with conventional gene transfer-based method. Among the mixture of the modified cells on the substrate, the intercellularly contacting cell was observed to be fluorescently labeled by confocal microscopy. Similarly, in cell suspensions, the intercellularly contacting cell of interest could be labeled and quantitatively analyzed by flow cytometry. In a variety of analytical systems for cell-cell communications, the present simple and versatile method for labeling the intercellularly contacting cells is a potentially promising tool in accurate distinguishment of the cellular responses due to intercellular contact.

## Acknowledgments

This research was supported by Japan Science and Technology Agency (JST) the MIRAI program 19217334. T. K. was supported by a Grant-in-Aid from the Japan Society for the Promotion of Science (JSPS) Research Fellows.

